# A genome-wide screen in *Pseudomonas aeruginosa* identifies genes impacting production of the hemolytic phospholipase C/sphingomyelinase, PlcH

**DOI:** 10.64898/2026.04.23.720442

**Authors:** Kristin Schutz, Olivia F. Evans, Jacob R. Mackinder, Pauline DiGianivittorio, Aditya Patwardhan, Matthew J. Wargo

## Abstract

The secreted phospholipase C/sphingomyelinase, PlcH, is the heat-labile hemolysin of *Pseudomonas aeruginosa* and one of its important secreted virulence factors. While there are known and suspected genes that impact PlcH production in *P. aeruginosa*, we sought to identify additional genes by screening the PA14 transposon mutant library to measure extracellular PlcH enzyme activity induced by choline. The library as a whole had a log2-normal distribution of NPPC activity with notable tails that included the genes of interest. These outlier genes included nearly all of those known to be important for PlcH production in response to choline, including those required for choline metabolism, glycine betaine sensing, and secretion through the outer membrane. Interestingly, higher PlcH production was also seen in mutants of the protease associated genes *lon*, *mucD*, and *clpA*, as well as other genes. Additionally, we identified genes impacting baseline levels of PlcH production, which include genes in the dimethylglycine metabolism locus involved in choline metabolism. The high hit rate of known and suspected genes supports the power of this screen and our verification of these genes by clean deletion in strain PA14 confirm the broad importance of these systems across *P. aeruginosa*, as previous work was confined to strain PAO1. There were many genes identified in this screen that were not individually examined and the complete screen results reported here should allow others to identify intersection of their genes of interest with PlcH production.

**Importance:** *Pseudomonas aeruginosa* is an important opportunistic pathogen that employs multiple independent virulence factors to cause infection, one of which is the hemolytic phospholipase C/sphingomyelinase PlcH. Using a whole genome screen, we identified both known and previously unknown genes contributing to *P. aeruginosa* PlcH production. Our findings provide insight into the integration of various cellular processes with PlcH production and identify potential genes that may impact the PlcH expression heterogeneity seen in *P. aeruginosa* clinical isolates.

## Introduction

*Pseudomonas aeruginosa* is a common opportunistic pathogen, particularly in healthcare settings, that causes infections in a variety of body sites including the skin, eyes, lungs, and bloodstream. Morbidity and mortality due to *P. aeruginosa* infections is driven in part by a collection of virulence factors secreted either directly into host cells by the type 3 and type 6 secretion systems or into the extracellular milieu via other secretion systems where they can directly impact host molecules and cells^1–3^. One of these extracellular secreted virulence factors is the heat-labile hemolysin^4^, PlcH, which is a secreted phospholipase C/sphingomyelinase^5^. PlcH expression is retained in all clinical strains tested^6–9^ and higher PlcH production by strains has been negatively correlated with patient outcomes and azithromycin efficacy in cystic fibrosis^9,10^. Mutants in *plcH* are less virulent in infection models^11–16^ and PlcH directly drives cytotoxicity, inflammation, and pulmonary surfactant damage^11,17–21^.

PlcH is secreted from the cytosol to the periplasm via the twin-arginine translocase (TAT) system^22^ and secreted from the periplasm to the extracellular milieu via the general secretion pore (Gsp, also called Xcp in *P. aeruginosa*)^23,24^. PlcH secretion is aided by the PlcR1 and PlcR2 chaperones, both encoded by the *plcR* gene in the *plcHR* operon^25^. Transcription of *plcH* can be induced by phosphate starvation via PhoB^26,27^, glycine betaine (GB) and dimethylglycine via GbdR^28^, and sphingosine via SphR^29^. Choline can induce *plcH* transcription in a *gbdR*-dependent manner, but only once it has been metabolized to GB, as mutations in the choline oxidase, *betA*, prevent *plcH* induction by choline^30^. Induction of *plcH* by sphingosine via SphR occurs due to transcriptional readthrough from the upstream *cerN* promoter^29^. Anr also regulates *plcH* transcription and this regulation is impacted by GB metabolism^31,32^. Transcription induction of *plcHR* by GB can be regulated by catabolite repression control^33^, likely due to catabolite repression of *gbdR* transcription^34^, though this has not been definitively tested and it is important to note that many clinical strains have mutations in or quickly lose catabolite repression during infection^35^. There is a Vfr binding site in the *cerN*-*plcH* intergenic region though no function due to this site has been previously shown and Vfr was previously ruled out as the mediator of *plcH* catabolite repression^33^. The H-NS family proteins MvaU and MvaT are known to regulate expression from many loci in *P. aeruginosa*, directly or indirectly, including *cerN* and *plcH* ^36^. PhrS has also been shown to negatively regulate *plcH* expression, though likely indirectly^37^.

Given these known and suspected genes impacting PlcH expression and the potential that other genes also contribute, we set out to identify transposon mutants with altered PlcH expression by screening the *P. aeruginosa* PA14 transposon mutant library for PlcH activity by measuring hydrolysis of the colorimetric substrate nitrophenylphosphorylcholine (NPPC). Here we report on the outcomes of that screen, further characterization of selected genes, and highlight some previously unsuspected genes that impact PlcH production in *P. aeruginosa*.

## Results

### Screen of the PA14 TnM library for altered choline-dependent induction of PlcH enzyme activity

We conducted nitrophenylphosphorylcholine (NPPC) hydrolysis assays on the entire PA14 mutant library^38^, as described in detail in the Materials & Methods, to identify mutants with altered extracellular PlcH enzyme activity. The NPPC assay is a sensitive proxy for PlcH activity, particularly using choline as the only inducing signal, as other NPPC hydrolyzing enzymes (particularly PlcN), are not induced under these conditions^27,30,33,39^. The within-plate normalized choline-induced NPPC hydrolysis activity shows a normal distribution when log2 transformed, with a log2 mean of 6.57 (95.2% linear), a median of 6.66 (101.1% linear), and standard deviation of 0.73 (**Figure 1**). The transposon mutants with the highest and lowest NPPC hydrolysis activities are listed with additional details in **Table 1**. Complete data for this screen is available in **Supplemental Table S1**, which includes Z-scores. Insertants that were determined to be auxotrophs or determined to be cross-contaminants by sequencing were removed from **Figure 1** and **Table1**. The two instances of cross-contamination are noted in text in **Supplemental Table S1**, while auxotrophs have “NA” listed in the activity column – described more in the Methods Section.

**Figure 1.**
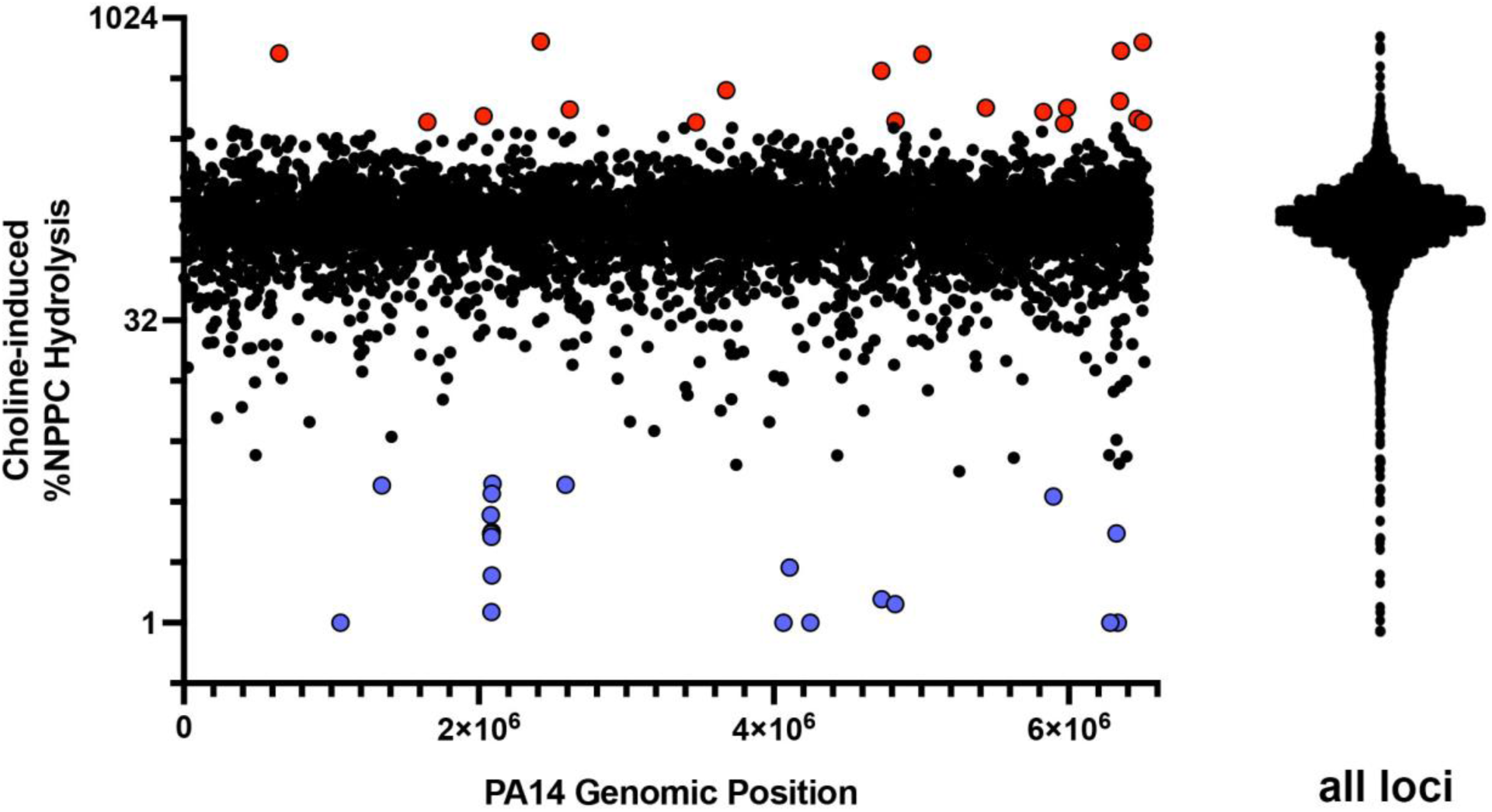
Screen of the *P. aeruginosa* PA14 transposon mutant library for choline-induced NPPC hydrolysis activity. The NPPC hydrolysis activities for each transposon mutant are plotted on a log2 scale after normalization to the within-plate average. On the left, expression is mapped versus the genomic position of each transposon insertion. Red points are those with the highest NPPC hydrolysis activity, while blue points are those showing the lowest NPPC hydrolysis activity, both classes of which are detailed in Table 1. To the right is the single column scatter plot of the same data showing the normal distribution of the transposon insertant activities.

**Table 1.**
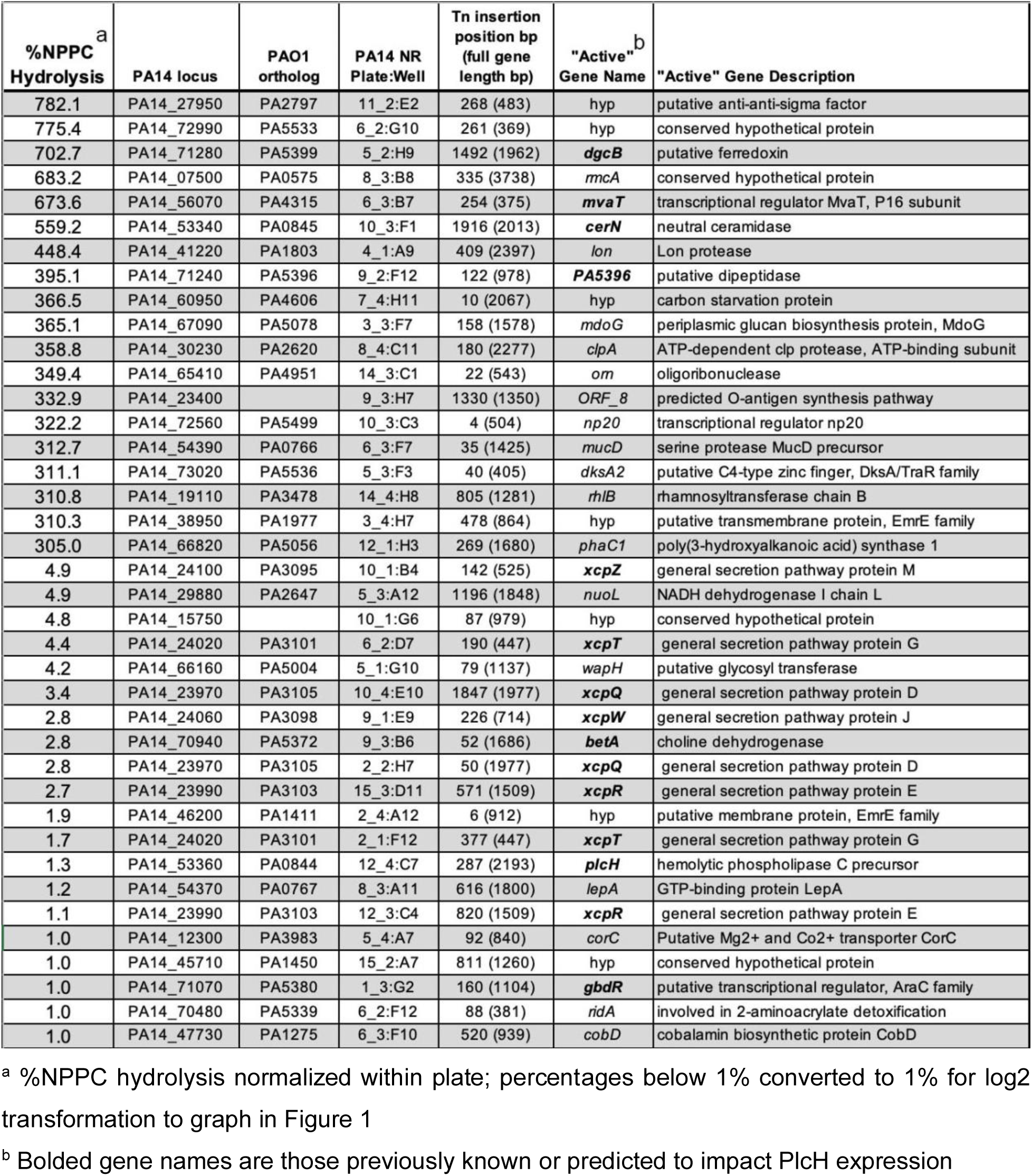
List of transposon mutants with the highest and lowest levels of choline-induced NPPC hydrolysis activity.

### Transposon insertions in known PlcH secretion and export systems, along with choline metabolism genes, validate the screen

Transposon insertion into genes previously known to impact PlcH expression provided internal validation for the screen. For the data in this paragraph, we are referring to **Figure 1**, **Table 1**, and **Supplemental Table 1**. The transposon insertion into *plcH* shows very low NPPC hydrolysis activity, supporting our NPPC hydrolysis assay as predominately measuring PlcH enzyme activity under these conditions. This conclusion is also supported by the insertion mutant in *gbdR*, which is required for choline-dependent induction of *plcH* transcription^28^. Transposon insertion in *betA* and, to a lesser extent *betB*, which encode the choline oxidase and betaine aldehyde dehydrogenase, respectively, result in reduced PlcH induction by choline, as expected^30^. We previously showed that chemical inhibition of the dimethylglycine catabolism enzyme, DgcA, or deletion of the *dgcA* gene in strain PAO1 led to hyper-induction of *plcH* transcription in the presence of choline^40^, and conversely, that over-expression of the GB catabolic genes *gbcA* & *gbcB* led to reduced PlcH expression^41^. Correspondingly, transposon insertions in the dimethylglycine catabolic operon (*dgcB* and *PA5396*) show the expected hyper-induction phenotype. At this timepoint in PA14, transposon insertions into *gbcA* and *gbcB* did not show hyperinduction of PlcH.

Transposon insertions into nearly every gene in the *xcp* locus were defective for extracellular PlcH activity, supporting previous findings that PlcH is exported from the periplasm by the Xcp/Gsp system^23^. The *xcp* transposon mutants also demonstrate that NPPC is not significantly transported into the periplasm in PA14, where PlcH remains active^23,24^. One system required for PlcH expression that was not identified in this screen was the TAT secretion system encoded by *tatABC*^42^. There are no insertants in *tatA* or *tatB* in the PA14 transposon library and the *tatC* insertion in our library copy did not show any of the known TAT phenotypes^42^. We determined that our *tatC* well was contaminated by a transposon mutant from an adjacent well and no *tatC* insertion was detectable by PCR from that well.

To verify the role of TAT and Xcp in PA14 PlcH secretion, as most prior work was conducted in *P. aeruginosa* strain PAO1 and the above data are from transposon mutants, we generated both *tatABC* and *xcpRSTUVWXY* deletion mutants in *P. aeruginosa* PA14 and measured extracellular NPPC hydrolysis activity compared to wild-type (**Figure 2**), which showed that both systems are required for functional extracellular PlcH, in concordance with results in PAO1^23,24,42^.

**Figure 2.**
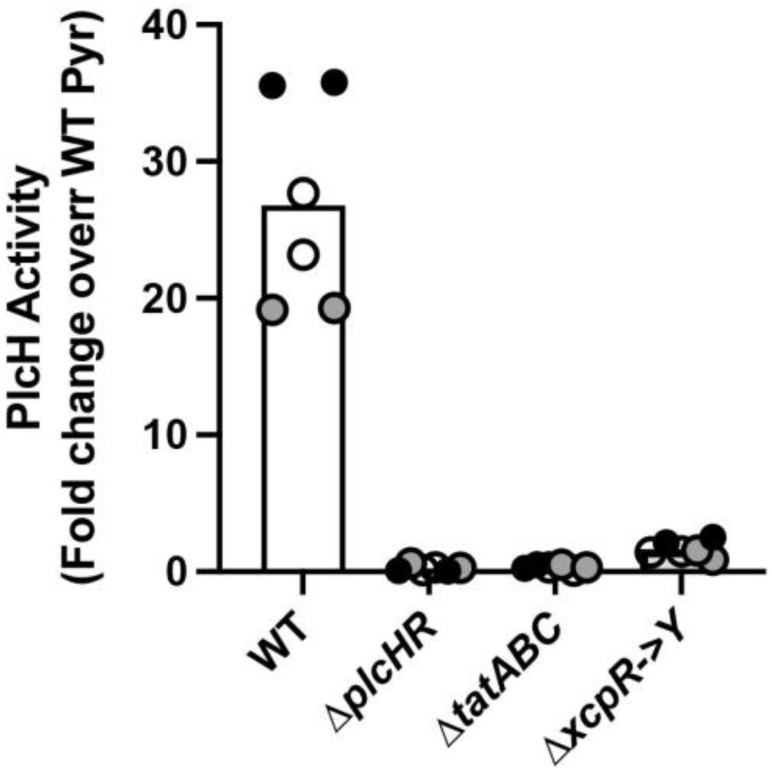
Secreted PlcH activity in mutants of the TAT secretion system and Xcp/Gsp system. The fold change in PlcH activity is plotted as measured by NPPC hydrolysis after choline induction over the hydrolysis in the non-inducing pyruvate control condition. The Δ*plcHR*, Δ*tatABC*, and Δ*xcpRSTUVWXY* (Δ*xcpR→Y*) strains are all statistically different than WT with p < 0.0001, determined by ANOVA with Dunnett’s post-test using WT as the comparator. All data points are shown and are colored by experiment with white circles for all replicates from experiment #1, grey from experiment #2, and black from experiment #3, with the overall means represented by the bars. Only the means from each experiment are used in the statistical analyses for these panels (i.e. n = 3 per condition).

### Secondary screen for transposon mutants not further explored in this study

We selected a subset of transposon mutants that were, for the most part, not further examined in this study and assessed PlcH activity induced by choline compared to induction by sphingosine. Since these two inducing conditions are dependent upon different regulators ^28,29^, they allow orthologous assessment of whether impacts are likely transcriptional or likely post-transcriptional. As seen in **Figure 3**, there is a rough log linear relationship between induction in each condition for most of these mutants, suggesting that their impact on PlcH expression is independent of the induction method (i.e. likely post-transcriptional). The two exceptions to this pattern to note are *mvaT::Tn* and *rhlB::Tn*, both of which show higher PlcH induction in choline than WT, but similar or lower induction in response to sphingosine. After confirmation of these phenotypes using the secondary screen, we explored a number of other genes identified in **Table 1** throughout the remainder of the Results section.

**Figure 3.**
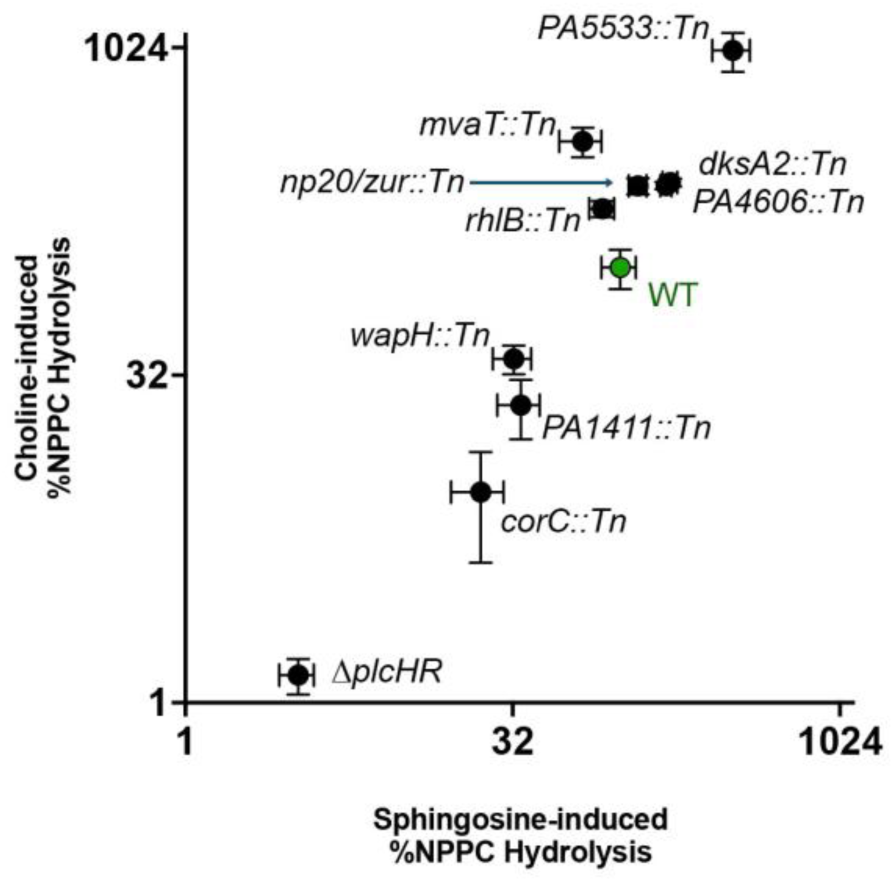
Secondary screen of selected transposon mutants for relative PlcH production. The NPPC hydrolysis activity induced by choline (y-axis) or sphingosine (x-axis) of a selection of transposon mutants was normalized to the levels of PA14 WT (in green). Error bars represent standard deviation of three independent biological replicates.

### Transposon insertions that lead to altered baseline NPPC hydrolysis activity

Most transposon mutants, including many of those with altered PlcH activity in response to choline, maintained normal NPPC hydrolysis activity in the un-induced condition (baseline), however a few showed marked induction at baseline (**Table 2; Supplemental Table 1**). The top transposon mutant altered for baseline PlcH production was an insertion into *cerN*. That particular insertion is at the extreme 3’ end of the *cerN* gene and our prediction is that the transposon internal promoter is constitutively driving baseline transcription of *plcH*, which lies immediately downstream of *cerN*. Deletion of *cerN* led to increased choline-dependent PlcH production, but that effect was not complemented in trans and the deletion did not have increased basal induction or increased induction in response to sphingosine (**Supplemental Figure 1**). Thus, the *cerN::Tn* baseline expression phenotype is likely dependent upon the position and presence of the transposon. However, there is a specific effect of the *cerN* deletion on choline-induced PlcH expression, but not sphingosine induction, suggesting some impact on GbdR function at the *plcH* promoter upon *cerN* deletion.

**Table 2.** Transposon mutants with more than 5-fold higher NPPC hydrolysis activity than WT in baseline (uninduced) conditions.

| Baseline (Pyr) %NPPC Hydrolysis Activity | Choline induced %NPPC Hydrolysis Activity | PA14 locus | PA01 ortholog | PA14 NR Plate:Well | Gene Name | Gene Description |
| --- | --- | --- | --- | --- | --- | --- |
| 30890.3 | 559.2 | PA14_53340 | PA0845 | 10_3:F1 | *cerN* | alkaline ceramidase |
| 6242.3 | 702.7 | PA14_71280 | PA5399 | 05_2:H9 | *dgcB* | ferredoxin in dimethylglycine catabolic operon |
| 1043.7 | 775.4 | PA14_72990 | PA5533 | 06_2:G10 |  | conserved hypothetical protein |
| 834.7 | 262.1 | PA14_70850 | PA5368 | 05_2:H12 | *pstC* | phosphate ABC transporter, permease protein |
| 711.6 | 118.1 | PA14_21090 | PA3320 | 03_1:D10 |  | conserved hypothetical protein |
| 708.2 | 142.4 | PA14_71460 | PA5415 | 01_2:E9 | *glyA1* | serine hydroxymethyltransferase |
| 639.0 | 191.4 | PA14_55820 | PA4297 | 08_3:H7 |  | putative membrane protein |
| 628.0 | 349.4 | PA14_65410 | PA4951 | 14_3:C1 | *orn* | oligoribonuclease |
| 610.6 | 683.2 | PA14_07500 | PA0575 | 08_3:B8 | *rmcA* | conserved hypothetical protein |
| 607.6 | 189.3 | PA14_58470 | PA4505 | 03_4:C8 | *dppD* | putative dipeptide ABC transporter |
| 587.1 | 322.2 | PA14_72560 | PA5499 | 10_3:C3 | *np20, zur* | transcriptional regulator np20 |
| 583.4 | 153.1 | PA14_58600 | PA4516 | 03_1:C3 |  | conserved hypothetical protein |
| 554.8 | 144.2 | PA14_43630 | PA1615 | 03_1:B7 |  | putative lipase |
| 521.7 | 97.9 | PA14_00730 | PA0061 | 05_4:A12 |  | conserved hypothetical protein |
| 504.0 | 234.0 | PA14_55060 | PA0987 | 06_4:B1 |  | conserved hypothetical protein |

Transposon insertion into *PA5533* led to increased baseline and induced levels of PlcH activity in response to both choline and sphingosine (**Table 1**, **Table 2** and **Figure 3**), but deletion of this gene did not alter PlcH activity (**Supplemental Figure 2**). The impact of transposon insertion into *PA5533* may be due to its proximity to the zinc regulator, *zur*, insertion into which showed a PlcH production phenotype (**Table 1**, **Figure 3**, **Table 2**, and **Supplemental Table 1**). The increased baseline phenotype of the *pstC* insertant is expected based on the constitutive activation of PhoB in this mutant, which is an activator of *plcH* transcription^27,43^. The increased baseline phenotype for the *dgcB* and *glyA1* insertants will be covered later in the Results section. We have not followed up on the other genes in **Table 2** at this time.

### Mutations in some protease-related genes enhance PlcH expression

Among our top mutants for PlcH over-production were mutants with transposon insertions in the *clpA*, *lon*, and *mucD* genes (**Table 1**), each of which encode a protease or component of a protease complex. We generated a clean deletion for each of these genes and, while generally phenocopying transposon insertants for overproduction (**Figure 4A**), we noted that for the *clpA* and *mucD* deletions the phenotype was dependent on both the media in the plate used for recovery from the -80°C and the multi-well plate well-size used during induction. Note, particularly, that Δ*mucD* shows trends in these experiments based on starting media and well-size (**Supplemental Figure 3**) but only its fold-induction in choline (**Figure 4B**) is statistically significant. The *lon* deletion shows over-production at baseline and over-production during induction, thus the fold induction between baseline and induced is not much higher than that of WT. Conversely, Δ*clpA* and Δ*mucD* both show PlcH activity in the uninduced condition similar to or lower than WT but a larger difference (fold induction) between uninduced and choline-induced conditions (**Figure 4B**) that varies based on the plate and induction conditions (data for all media and well combinations is shown in **Supplemental Figure 3**).

**Figure 4.**
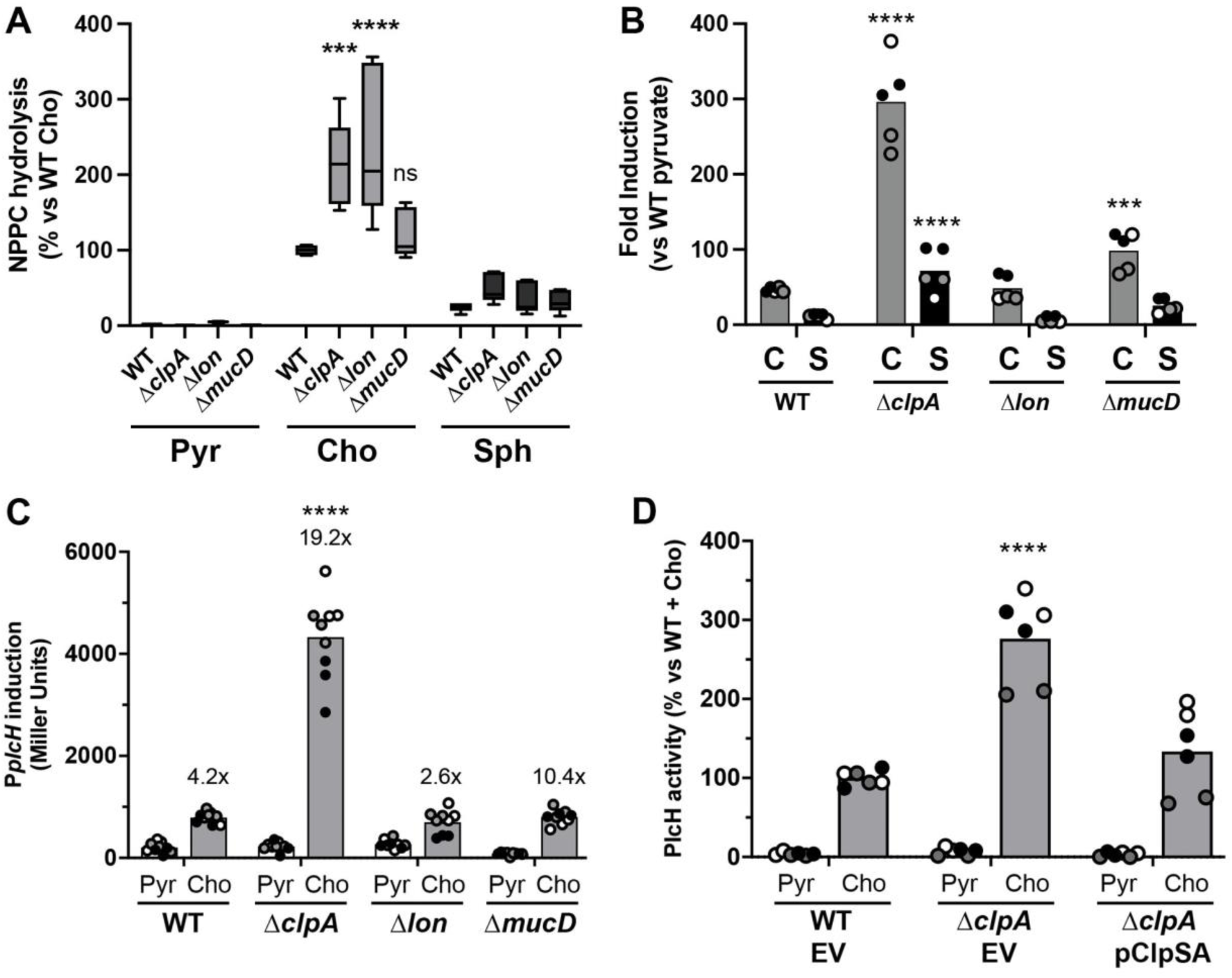
Impact of protease-related genes on PlcH activity. **(A)** PlcH activity of WT and mutants of *clpA*, *lon*, *mucD* in uninduced (pyruvate, Pyr) compared to induction with choline (Cho), or sphingosine (Sph). NPPC hydrolysis activity normalized to WT + choline as 100%. **(B)** Fold induction of NPPC hydrolysis activity based on the activity data presented in (A)**. (C)** Transcriptional reporter assay showing expression from the pMW22 P*_plcH_-lacZ* reporter plasmid comparing pyruvate and choline induction in each strain. The numbers over the bars for each strain denote the fold-change between pyruvate and choline conditions for this transcriptional reporter**. (D)** Complementation of NPPC hydrolysis activity with *clpSA* expression. Statistical analysis for all by two-way ANOVA with Dunnett’s post-test within each induction condition with WT as the comparator strain (***, p<0.001; ****, p<0.0001). All data points are shown and are colored by experiment with white circles for all replicates from experiment #1, grey from experiment #2, and black from experiment #3, and the overall means are represented by the bars. Only the means from each experiment are used in the statistical analyses for these panels (i.e. n = 3 per condition).

To determine whether PlcH over-production in these strains occurred at the level of choline-dependent transcriptional induction, we used our previously described P*_plcH_*-*lacZ* reporter plasmid, pMW22^28^. Deletion of *clpA* led to an increase in reporter activity from the *plcH* reporter compared to WT (**Figure 4C**). Deletion of *lon* and *mucD* did not increase total *plcH* reporter expression during induction with choline, but the *lon* deletion mutant showed higher promoter activity in the uninduced condition (and thus lower fold-induction), while the *mucD* deletion mutant showed lower promoter activity in the uninduced condition (thus higher fold-induction) (**Figure 4C**).

We have had long-term interest in *plcH* transcriptional regulation and thus wanted to further verify the impact of *clpA* mutation on PlcH expression. In addition to the *clpA* transposon mutant in **Table 1**, there is an additional *clpA* insertant in the library (1_2:B6) that showed ∼260% PlcH production (**Supplemental Table 1**). These two independent transposon mutants phenocopy a *clpA* deletion mutant, and a plasmid containing the *clpSA* operon complemented the PlcH over-production phenotype (**Figure 4D**).

### PlcH secretion in *plcR* mutants

Transposon insertants in the *plcR* gene failed to make our cutoff for **Table 1**, even though *plcR* deletion mutants were previously shown to have markedly reduced supernatant PlcH activity in PAO1^14,25^. Further examination of the insertion sites for the two *plcR* transposon mutants in the PA14 library (**Supplementary Table 1**) show that both have insertions prior to the start of the second in-frame open reading frame, termed *plcR2*. Thus, both transposon insertions disrupt the *plcR1* product but do not insert into the *plcR2* product, and *plcR2* is sufficient for PlcH secretion^25^. Since the work on *plcR* and initial investigation into the roles of *plcR1* and *plcR2* were previously conducted only in PAO1, we generated a *plcR* deletion mutant in PA14 and tested complementation with the empty vector, *plcR (plcR1 & plcR2),* or *plcR2* alone. In **Figure 5**, we show that both the *plcR* and *plcR2* genes show a similar function in PA14 as they do in PAO1. While reports in PAO1 show about a 90% reduction in NPPC hydrolysis activity in Δ*plcR* supernatants compared to WT, here we show that the PA14 Δ*plcR* strain has about a 60% reduction in NPPC hydrolysis activity in the supernatant. These differences may be partly due to media and induction time differences between the studies, but could certainly be strain-dependent differences in importance of PlcR in PlcH folding, secretion, and/or activity.

**Figure 5.**
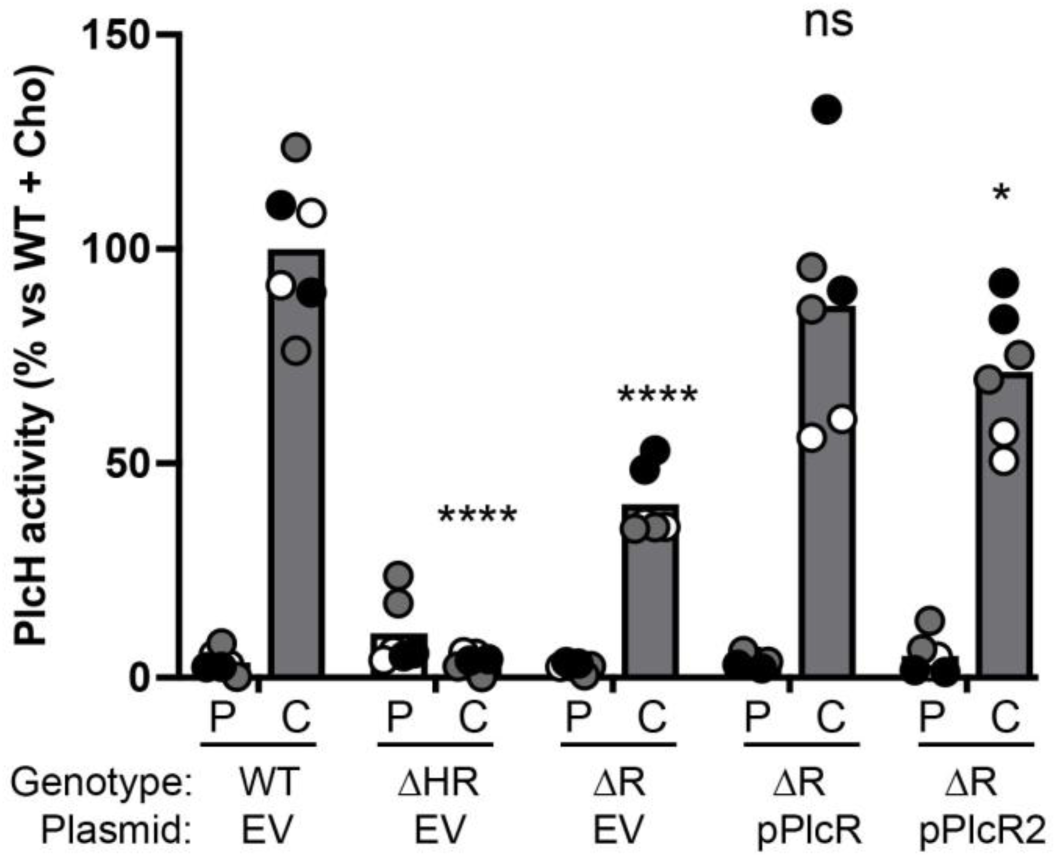
The impact of *plcR* on PlcH activity in PA14. Normalized NPPC hydrolysis activity from *P. aeruginosa* PA14 WT, Δ*plcHR* (ΔHR), Δ*plcR* (ΔR) carrying the pUCP22 vector (EV), the full *plcR* gene encoding both PlcR1 and PlcR2 (pPlcR), or only the *plcR2* gene (pPlcR2). Statistical analysis by two-way ANOVA with Dunnett’s post-test within each condition with WT as the comparator strain (****, p<0.0001). All data points are shown and are colored by experiment with white circles for all replicates from experiment #1, grey from experiment #2, and black from experiment #3, and the overall means are represented by the bars. Only the means from each experiment are used in the statistical analyses for these panels (i.e. n = 3 per condition).

### The impact of mutations in the choline catabolic pathway on PlcH production

Two transposon insertants with both increased baseline and choline-induced PlcH activity were in the *dgcB* and *glyA1* genes that are part of the choline catabolic pathway^44^. The impact of these mutations during choline-dependent induction was anticipated^45^, but the increase in PlcH baseline production was unexpected. To follow up on this observation, we measured PlcH production in uninduced and choline-induced conditions comparing WT to deletion mutants in genes for each step of the *P. aeruginosa* choline catabolic pathway (**Figures 6A & B**). In the choline-induced condition, deletion of *dgcA*, *dgcB*, or the entire *dgc* operon (*PA5396*, *PA5397*, *dgcA*, & *dgcB*) resulted in increased NPPC hydrolysis activity (**Figure 6A**). In the baseline condition, these three mutants were also the only ones to show significant increase in activity (**Figure 6B**), though there was much more variability in the basal phenotype than the induced phenotype, and deletion of *dgcA* had a greater impact than deletion of *dgcB*. The reason for increased basal activity in mutants of the *dgc* locus is not known, though we speculate in the discussion.

**Figure 6.**
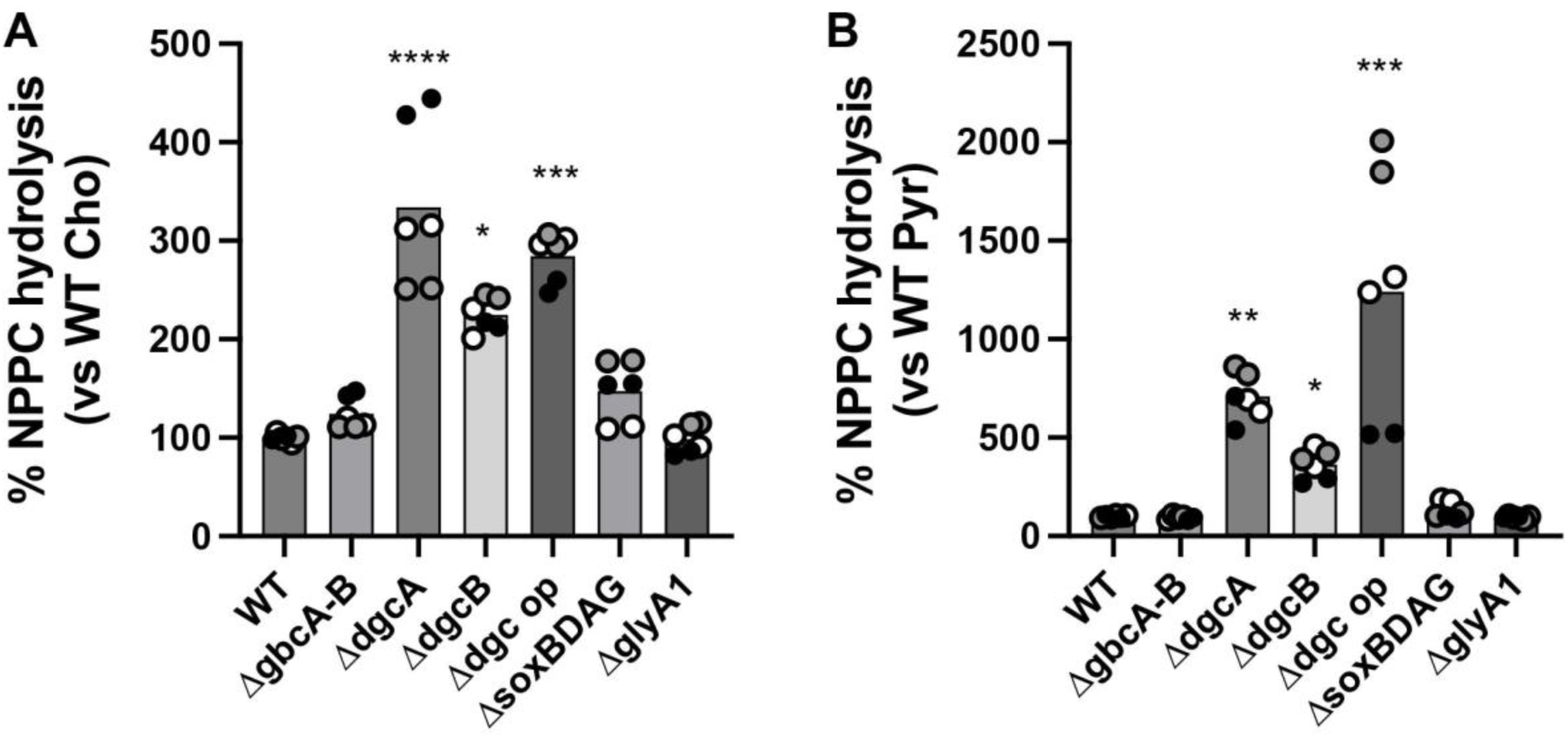
PlcH activity is altered for some mutants in the choline catabolic pathway. NPPC hydrolysis activity measured **(A)** under choline induction conditions or **(B)** at baseline pyruvate conditions. Statistical analysis for each panel by ANOVA followed by Dunnett’s post-test with WT as the comparator strain (*, p<0.05; **, p<0.01; ***, p<0.001; ****, p<0.0001). All data points are shown and are colored by experiment with white circles for all replicates from experiment #1, grey from experiment #2, and black from experiment #3, and the overall means are represented by the bars. Only the means from each experiment are used in the statistical analyses for these panels (i.e. n = 3 per condition).

Interestingly, while the *glyA1* transposon insertion mutant showed increased PlcH activity at baseline (**Table 2**), we did not see that in either the Δ*glyA1* or the Δ*soxBDAG* strains (**Figure 6B**). The *glyA1* gene is the first gene in the *glyA1soxBDAG* operon, so perhaps a deletion of the whole operon would show high baseline PlcH activity, mimicking the *glyA1* transposon mutant, though we have not tested that hypothesis.

### Further examination of two additional genes identified in this screen

Transposon insertion into *rmcA* led to both increased baseline and choline-induced PlcH production (**Tables 1 & 2**, **Supplemental Table 1**). The *rmcA* gene is a multifunctional protein containing the cyclic-di-GMP-related domains GGDEF and EAL as well as a redox-related PAS domain and is involved in *P. aeruginosa* biofilm formation^46^. We received the Δ*rmcA*, WT parent, and *attTn7* complementation strains from George O’Toole (Geisel School of Medicine at Dartmouth) and measured PlcH production at baseline and during induction with choline (**Figure 7**). The Δ*rmcA* strain shows neither increased baseline nor increased choline induced NPPC hydrolysis activity, but the complemented strain does show a significantly higher level of PlcH activity during choline induction. To better understand the discrepancy between the transposon mutant observation and the clean deletion, we examined the position of the transposon insertion. The *rmcA* transposon insertant that made our PlcH hyperproduction list was 8_3:B8 with the transposon inserting at base 335 of 3738. There is a second transposon insertant in the PA14 library, 15_3:F11 (insertion at base 3425 of 3738), that has no significant PlcH expression phenotype. The 15_3:F11 insertion cuts off the C-terminal EAL domain, whereas the 8_3:B8 insertion falls in the periplasmic domain. Potentially, expression of the remaining portion of the *rmcA* coding sequence from an internal start codon leads to a situation where the allele is over - expressed and/or has altered regulatory function.

**Fig. 7.**
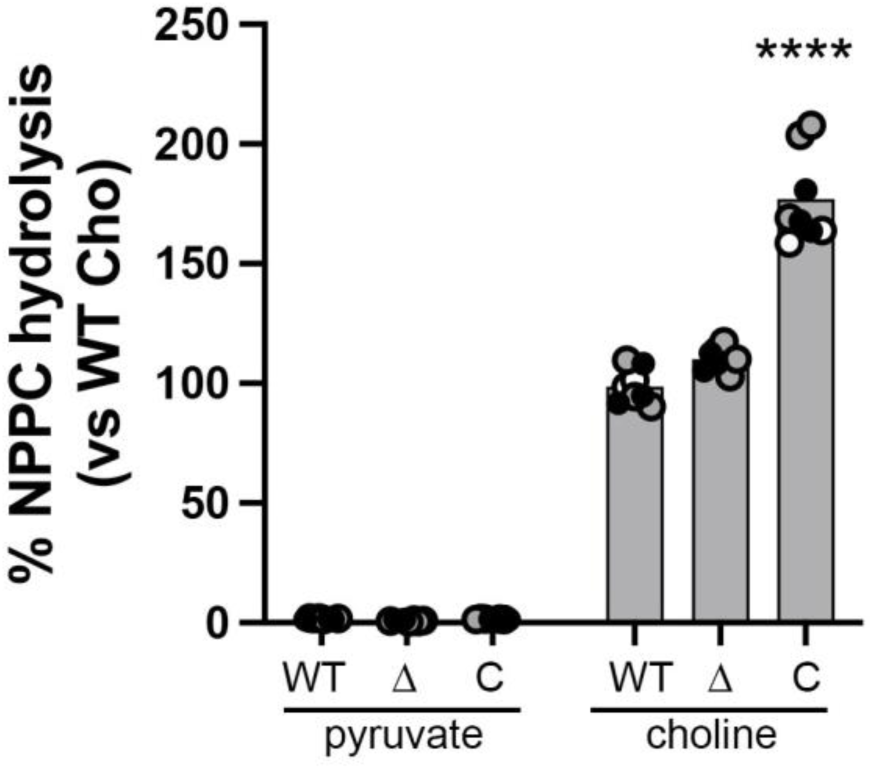
Overexpression of *rmcA* increases PlcH expression. PlcH activity as measured by NPPC hydrolysis comparing WT, Δ*rmcA* (Δ), and the Δ*rmcA attTn7::rmcA* complemented strain (C) in pyruvate and choline. Statistical analysis by two-way ANOVA with Dunnett’s post-test within each condition with WT as the comparator (****, p<0.0001). All data points are shown and are colored by experiment with white circles for all replicates from experiment #1, grey from experiment #2, and black from experiment #3, and the overall means are represented by the bars. Only the means from each experiment are used in the statistical analyses for these panels (i.e. n = 3 per condition).

*PA5339* encodes RidA, loss of which impacts growth and motility via accumulation of 2-aminoacrylate^47^. In our initial screen, the *PA5339::Tn* strain had a low PlcH activity phenotype (**Table 1**). We deleted the *ridA* gene and noticed immediately that it did not grow well in our standard overnight culture conditions (MOPS pyruvate + glucose). The *ridA* deletion also had growth phenotypes on choline, glycine betaine, and dimethylglycine, but not in MOPS with 0.05% tryptone (**Figure 8A**). When we grew overnight cultures in the MOPS + 0.05% tryptone and then transferred to standard induction conditions, we noted increased NPPC hydrolysis in the Δ*ridA* mutant in the MOPS pyruvate + choline condition compared to WT (**Figure 8B**). This suggests that the initial screen phenotype was likely impacted by limited cell growth, but that changing the pre-growth conditions uncovered over-production of PlcH in the *ridA* mutant based on its inability to metabolize choline.

**Figure 8.**
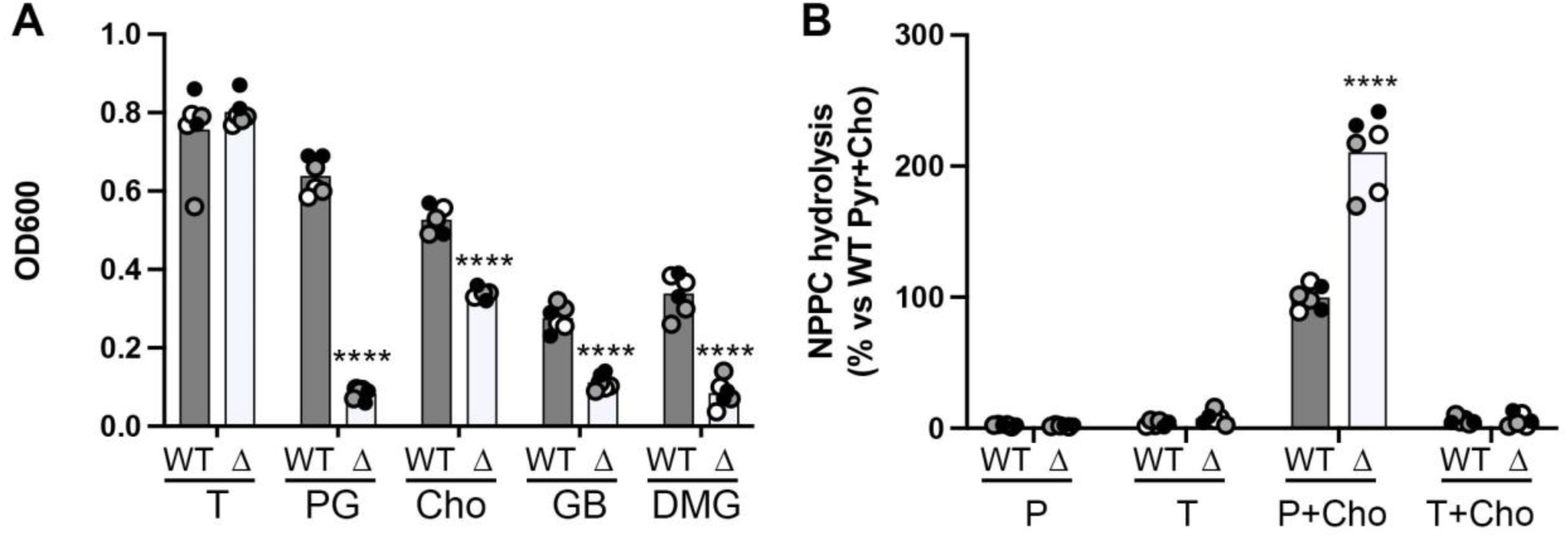
Deletion of *ridA* (*PA5339*) leads to increased PlcH expression correlated with growth defects on choline and its metabolites. (A) Growth of WT vs Δ*ridA* (Δ) at 24 hours as measured by OD_600_ from cells started at OD_600_ = 0.05. MOPS media with no carbon source was the base and the added carbon sources are noted: 0.05% tryptone broth (T), 20 mM pyruvate + 5 mM glucose (PG), 20 mM choline (Cho), 20 mM glycine betaine (GB), or 20 mM dimethylglycine (DMG). **(B)** PlcH activity measured as NPPC hydrolysis from cells grown as overnights in MOPS + 0.05% tryptone broth and moved to the listed conditions for induction. MOPS media with no carbon was the base media and the added compounds are noted: 20 mM pyruvate (P), 0.05% tryptone broth (T), 20 mM pyruvate + 2 mM choline (P+Cho), or 0.05% tryptone broth + 2 mM choline (T+Cho). Statistical analysis for both panels done with two-way ANOVA with Sidak’s post-test comparing WT to the *ridA* deletion strain in each condition (****, p<0.0001). All data points are shown and are colored by experiment with white circles for all replicates from experiment #1, grey from experiment #2, and black from experiment #3, and the overall means are represented by the bars. Only the means from each experiment are used in the statistical analyses for these panels (i.e. n = 3 per condition).

## Discussion

PlcH is an important virulence factor in *P. aeruginosa* and while focused investigations have uncovered more about its function and regulation recently, here we chose to explore PlcH regulation by screening the entire PA14 transposon mutant library for PlcH enzymatic activity. In doing so, we identified a number of genes in which insertion alters PlcH activity – some known, some suspected, and some unknown. We chose to follow up on a subset of these genes and present new connections between these pathways and PlcH.

For the most part, the impact of mutations in the choline pathway on PlcH activity were predicted based on previous results. What was not predicted was the much higher baseline activity of mutants in the genes of the *dgc* operon (*PA5396-PA5397-dgcA-dgcB*). In our examination of the GbdR regulon^45^, we only compared the *dgcA* mutant to WT in the choline condition, where hyperinduction of *plcH* and other GbdR-regulated transcripts was noted. Here, we observed the high basal levels of PlcH activity in these mutants (**Figure 6**). This suggests two possibilities. First, even though the strains are picked into media with no choline from the plate and grown overnight, it is possible that the population or some proportion of cells in the population retains enough choline from the plates to produce the observed induction. The second possibility is that there is synthesis of dimethylglycine or some related metabolite sensed by GbdR at a very low baseline level in *P. aeruginosa* that is normally removed by action of the Dgc enzymes, but builds up in their absence. We continue to examine both of these possibilities.

In addition to the known and suspected pathways identified, we were surprised by the phenotypes for *clpA*, *mucD*, and *lon* mutants (**Figure 4**). The mutants these genes encoding proteases (MucD and Lon) or the protease-associated chaperone (ClpA) showed increased PlcH activity which, for *mucD* and *clpA* mutants, was dependent upon growth conditions (**Supplemental Figure 3**). The function of Lon appears to be the most straightforward. Loss of *lon* results in increased baseline and induced PlcH activity (**Figure 4A**) and, while there seems to be a small impact on transcription in the lon mutant (**Figure 4C**), we suspect that PlcH is likely a substrate for Lon. Conversely, loss of *clpA* results in greater induction of both PlcH enzyme activity (**Figure 4A-B**) and transcriptional induction of *plcH* (**Figure 4C**). Though tempting to implicate regulation through the choline metabolic pathway, the *clpA* mutant also shows much higher PlcH activity than WT in the presence of sphingosine (**Figure 4B**), suggesting a broader impact of ClpA on *plcH* transcription. To date, our data do not resolve how much of the *mucD* phenotype is dependent on alteration of *plcH* transcription versus MucD directly targeting PlcH for degradation in the periplasm.

The *clpA* gene is retained in all Pseudomonads but, surprisingly, is frequently mutated during long-term *P. aeruginosa* infection in lungs of people with cystic fibrosis^48–51^. During in vitro evolution for growth with ceftazidime, *clpA* mutants are selected in both standard laboratory strains and hypermutator lineages^52–54^ and *clpA* mutants also arise in vitro during growth in aztreonam^51^. These in vivo and in vitro mutations are predominately in conserved residues or result in frameshifts, suggesting that many of these alleles are loss of function. Experimentally, transposon insertions in *clpA* result in lower virulence in *C. elegans* ^55^ but, conversely, increased survival in the murine lung ^56^. These findings all point to ClpA playing an important and probably multifaceted role in the CF lung and in response to ceftazidime and aztreonam. Here we demonstrate that *clpA* deletion increases *plcH* transcription and resultant secreted PlcH activity. It remains to be tested whether the *clpA* mutants arising in CF and in these in vitro selections always result in PlcH over-expression.

One of the weaknesses of this screen is that its cost and time-intensive enzymatic assay meant that the whole library could only be screened once. Thus, there are certainly false positives and false negatives present in this data that replication could have eliminated. As a discovery process, we were not particularly worried about false negatives, especially given the number of mutants with interesting phenotypes. Based on the genes previously known to have altered PlcH activity that had transposon insertions in the library, we hit 23 of 24, though not all were robust enough to be included in **Table 1**, suggesting a rough estimate of 4% for a false negative rate. False positives are more problematic in such a screen, and it is important to assess false positives that were methodological or stochastic versus false positives based on the nature of the transposon insertion site. Of the transposon mutants we screened secondarily (**Figure 3**) or tested with clean deletions, we had no transposon mutants whose phenotypes failed to repeat. We did, however, have a number of transposon mutant phenotypes that were not matched by clean deletion, but in which the gene had some role in PlcH activity (*rmcA*, *cerN*, *ridA*) and one with no phenotype at all (*PA5533*). In addition, we tried to make a deletion mutant for *PA2797* multiple times, but of dozens of double crossover colonies screened from three independent single crossovers, all were WT revertants.

We were not the first group to screen for mutants in PlcH activity. In 1997, Mike Vasil and colleagues reported results from a screen of ∼7500 random Tn5 transposon mutants for altered PlcH enzyme activity^30^. They had two hits that passed the secondary screen: *betB*, which encodes the betaine aldehyde dehydrogenase and *truB* (*PA4742*, *PA14_62730*), which encodes the tRNA pseudouridine 55 synthase. Transposon mutants for both of these are present in the PA14 library, and we covered *betB* along with other genes involved in choline metabolism in the results section. In our screen, the *truB* insertant showed ∼55% of normalized NPPC hydrolysis activity. Interestingly, *truB* was not the only tRNA modification-related gene identified as more than 30% up or down in our screen. The *PA4852* gene encodes a putative tRNA dihydrouridine synthase and showed 37% of normalized PlcH activity, while *PA3626*, encoding a putative tRNA pseudouridine synthase D, showed 55% of normalized PlcH activity. Transposon insertion into *trmA* (*PA4720*), encoding the tRNA uracil 5 methyltransferase, showed 132% PlcH activity. Additionally, while *trmD* is essential, the transposon mutant in the library has the insertion at base 757 of 759 and this strain shows 196% PlcH activity. In their paper, Vasil and colleagues noted that a *truB* mutant had more abundant *plcH* transcript but very low PlcH enzyme activity^30^. Along with *truB*, the other mutants in tRNA modification genes suggests there may be some translational regulation of PlcH expression that remains unexplored.

As with any genome-wide screen, there remains a lot of interesting genes that we have not followed up with additional experiments. One area of interest are genes related to zinc, including the >300% PlcH activity levels of mutants in the zinc related genes *zur* (*np20*) and *dksA2* (**Table 1**, **Figure 3**). In addition to these two large effects, insertion into the periplasmic binding protein of the Znu ABC transporter, *PA5498*, resulted in 150% PlcH activity, while insertion into *znuC* (*PA5500*) resulted in 120% PlcH activity. At this time, we do not know if the link to zinc and the signal of zinc starvation works directly on PlcH at some level or is mediated by affecting choline metabolism, cofactors of which require zinc, but something we are interested in exploring. Our findings in this screen and its follow-up experiments cement the role of known and suspected genes, providing more detail on their impact and verifying function in the *P. aeruginosa* strain PA14. Beyond those, the screen also identified a number of genes that impact PlcH activity that were unexpected. Some of these were characterized here, but the larger proportion remain unexplored. Likely others will be able to discover connections between their genes of interest and production of this important *P. aeruginosa* virulence factor using the data presented here.

## Materials and Methods

### Strains and growth conditions

Strains were stored as glycerol stocks at -70 or -80°C and recovered on Pseudomonas Isolation Agar (PIA) or PIA with 50 µg/ml gentamicin as appropriate for the strain. Specific instances of recovery on LB from freezer stocks is noted in the results section. During genetic manipulations, *P. aeruginosa* was selected for, and *E. coli* eliminated, using PIA plates supplemented with 50 µg/mL gentamicin. Prior to PlcH induction studies, *P. aeruginosa* was grown overnight in morpholinepropanesulfonic acid (MOPS) medium^57^ modified as previously described^58^, and supplemented with 20 mM pyruvate, 5 mM glucose, and 25 µg/ml gentamicin when appropriate.

### Generation of deletion mutants and complementation constructs

All PCRs were conducted using Q5 DNA polymerase (NEB). Primer sequences are listed in **Supplemental Table 2**. All plasmids created for this study were sequenced using Plasmidsaurus (Eugene, Oregon, USA). For plasmid carriage, *P. aeruginosa* and *E. coli* strains were transformed via electroporation and selected for growth on PIA with 50 µg/mL gentamicin and LB with 10 µg/mL gentamicin, respectively.

The allelic exchange vectors for the deletions generated in this study in **Table 3** were built using HiFi assembly (NEB) between a synthetic DNA block (IDT or Twist) and HindIII and KpnI digested pMQ30. After transformation into DH5a, miniprep, and sequencing, clones with correct and identical sequences were pooled for transformation into the conjugative donor *E. coli* strains S17λpir. Plasmid sequences for each of the deletion constructs are available in **Supplemental Data File 1**. Donor *E. coli* were mixed with their appropriate recipient strain and conjugation allowed to occur in spots on LB plates at 30°C overnight. Conjugation spots were resuspended and plated on PIA with 50 µg/ml gentamicin to select for single-crossover integrants. After restreaking onto PIA with 50 µg/ml gentamicin, single-crossover integrants were moved to LB without selection for 3 h prior to plating on LB with 5% sucrose and without salt at 30°C overnight. Sucrose resistant colonies were screened for loss of gentamicin resistance prior to PCR screening to determine whether each double-crossover colony was a mutant or WT revertant. PCR primers sequences for screening of each deletion mutant are listed in **Supplementary Table 2**.

**Table 3:** Strains and plasmids used in this study.

| **Strains:** |  |  |
| --- | --- | --- |
| **database #** | **Genotype** | **Source** |
| MJ1164 | PA14 WT | PMID: 7604262 |
| MJ1201 | $\Delta plcHR$ (restock of MJ144) | PMID: 38411048 |
| MJ1225 | $\Delta plcR$ | this study |
| MJ1059 | $\Delta tatABC$ | this study |
| MJ1272 | $\Delta xcpRSTUVMXY$ | this study |
| MJ1220 | $\Delta lon$ | this study |
| MJ1222 | $\Delta mucD$ | this study |
| MJ1138 | $\Delta clpA$ | this study |
| MJ587 | $\Delta cerN$ | PMID: 24465209 |
| PD146 | PA14 WT *attTn7::empty* | this study |
| PD148 | $\Delta cerN$ *attTn7::empty* | this study |
| PD150 | $\Delta cerN$ *attTn7::cerN* | this study |
| GO232 | PA14 WT | PMID: 33531388 |
| GO6279 | $\Delta rmcA$ | PMID: 33531388 |
| GO8813 | $\Delta rmcA$ *attTn7::PBAD-rmcA* | PMID: 33531388 |
| MJ288 | $\Delta gbcA-B$ | PMID: 17951379 |
| MJ336 | $\Delta dgcA$ | PMID: 17951379 |
| MJ1211 | $\Delta dgcB$ | this study |
| MJ1214 | $\Delta dgc$ operon (*PA5396-dgcB*) | this study |
| MJ1302 | $\Delta glyA1$ | this study |
| MJ1305 | $\Delta soxBDAG$ | this study |
| MJ1237 | $\Delta ridA$ (*PA5339*) | this study |
| MJ1295 | $\Delta PA5533$ | this study |
| **Plasmids:** |  |  |
| **database #** | **Desc** | **Source** |
| pUCP22 | empty expression vector | PMID: 1899844 |
| pMQ30 | allelic exchange vector | PMID: 16820502 |
| pTNS3 | Tn7 transposase expression | PMID: 18156318 |
| pFLP2 | Flp recombinase expression | PMID: 9661666 |
| pMW22 | *PplcH-lacZYA* reporter plasmid | PMID: 19103776 |
| pMW247 | pPlcR2 complementation vector | this study |
| pMW258 | pPlcR complementation vector | this study |
| pMW266 | pClpSA complementation vector | this study |
| pPD42 | *cerN* complementation via *attTn7* | this study |

For all but the *cerN attTn7* complementation vector (pPD42), complementation constructs were generated by HiFi of synthetic DNA fragments (Twist) into HindIII and KpnI cut pUCP22, putting each construct under control of its native promoter in opposite orientation to the transcription of the *lacZ* alpha fragment. Plasmid sequences for each complementation vector are available at **Supplemental Data File 1**.

The plasmid enabling chromosomal *attTn7*::*cerN* integration was generated by amplification from PA14 genomic DNA with primers #2869 and #2870 to generate a 2420 bp PCR fragment that was digested with HindIII and ligated into HindIII-cut pUC18-mini-Tn7T (CITE). After transformation into *E. coli* and miniprep, orientation was assessed by BamH1 digest and clones with proper orientation were sequenced. The pPD42 sequence is available at **Supplemental Data File 1.** To generate the chromosomal complementation and control strains, Δ*cerN* mutant and PA14 WT were co-electrotransformed with either pPD42 or pUC18T-miniTn7T-Gm along with pTNS3 based on the methods of Schweizer and colleagues^59^. Integrants at the *attTn7* locus were selected by growth on gentamicin and insertion verified by PCR. The gentamicin resistance cassette was excised by FLP-mediated recombination by electrotransforming the target strains with pFLP2, selecting on carbenicillin, and subsequently screening for loss of gentamicin resistance^59^. Loss of the gentamicin resistance gene was verified by PCR with primers #2771 and #2772 (**Supplemental Table 2**).

### Screen of the PA14 transposon mutant library for NPPC hydrolysis activity

To screen the PA14 transposon mutant library^38^, between two and four library plates per day were replicated onto LB plates with 10 µg/ml gentamicin using a multipin device and incubated overnight at 37°C. These overnight plates were replicated into 96-well polystyrene plates containing 200 µl/well of MOPS media with 20 mM pyruvate, 5 mM glucose, and 5 µg/ml gentamicin using a multipin device. The 96-well plated were sealed with a sterile, gas-permeable membrane (Microporous Sealing Film, USA Scientific) and incubated with horizontal shaking at 37°C for 18 hours. Auxotrophs were defined as having an increase in OD_600_ of less than 0.05 in the overnight growth plate.

From each column of the overnight plates, 50 µl was moved to two separate columns of a new plate with 150 ul of MOPS media, one with 20 mM pyruvate and the other with 20 mM pyruvate + 2 mM choline. Thus, each well from the overnight growth 96-well dish was moved to a non-inducing condition well (no choline) and an inducing condition well (2 mM choline), with each initial plate now distributed over two 96-well plates. These induction plates were sealed with a gas-permeable membrane and incubated at 37°C with horizontal shaking for three hours. Mutichannel pipetting was used to move 75 µl from each well into a fresh 96-well plate. After the plate was filled, 75 µl of 2x NPPC reaction buffer (based upon the Kurioka and Matsuda method ^60^, modified as we have described previously ^28^, but using a final concentration of 10 mM NPPC) was added with mixing by gentle pipetting and plates moved immediately to a pre-warmed (37°C) plate reader where, after the initial OD_600_ reading, plates were read at 410 nm every five minutes for 30 minutes. To determine the percent induction for NPPC hydrolysis activity, the change in 410 nm absorbance over time in the baseline (uninduced) condition was subtracted from the change in 410 nm absorbance over time in the induced (2 mM choline) condition. This delta-NPPC number was divided by the average delta-NPPC for that plate’s induced wells, not including auxotrophs and the well itself. Thus, percent NPPC hydrolysis is within-plate and within-day normalized for each well. The resulting log2 normal distribution (**Figure 1**) supports that this was a reasonable normalization strategy.

To determine the percent baseline NPPC hydrolysis, the change in 410 nm absorbance over time in the baseline (uninduced) condition was calculated and divided by the average delta-NPPC for that plate’s uninduced wells, not including auxotrophs and the well itself. Since the NPPC hydrolysis level is very low in WT and most of the transposon mutants, we focused only on mutants in which NPPC hydrolysis activity was increased.

### Induction and measurement of PlcH activity following the initial screen

For standard PlcH enzymatic assays, *P. aeruginosa* strains were grown overnight in MOPS minimal media with 20 mM sodium pyruvate, 5 mM glucose with 20 µg/ml gentamicin if appropriate. Cells were collected via centrifugation, washed in MOPS minimal media, and resuspended in MOPS media with 20 mM pyruvate or in MOPS with 20 mM pyruvate plus 2 mM choline or 50 µM sphingosine (Cayman Chemicals). All inductions took place at 37°C with horizontal shaking at 170 rpm for three hours, with plates covered by breathable films.

Phospholipase C activity was measured by observing the hydrolysis of the synthetic substrate ρ-nitrophenyl-phosphorylcholine (NPPC) based upon the Kurioka and Matsuda method ^60^, modified as we have described previously ^28^, but using a final concentration of 10 mM NPPC. NPPC hydrolysis was measured by monitoring the absorbance at 410 nm over time. Phospholipase C activity was calculated as micromoles of ρ-nitrophenol generated per minute of reaction per optical density (OD_600_). ρ-nitrophenol concentration was calculated using the extinction coefficient of 17700 M^-1^ cm^-1^ ^61^. Most of the PlcH activity data presented herein is normalized to the intraday activity of WT in either baseline or induced conditions, as noted in the results section and figure legends.

### Assessment of transcription induction using a *plcH-lacZ* reporter

To assess changes to transcription from the *plcH* promoter, we conducted β-galactosidase assays using a plasmid-based P*_plcH_-lacZ* transcriptional fusion as previously described^28^, using Miller’s method^62^. Induction conditions were identical to those described above for PlcH enzyme activity. Data were reported as Miller Units and fold induction was calculated relative to each strains’ uninduced (pyruvate) condition.

### Graphing and statistical analyses

All graphs were generated in GraphPad Prism after normalization and other relevant calculations in Microsoft Excel. Statistical analyses were based on one-way or two-way ANOVA with appropriate post-tests. We focused on differences of mutants compared to WT in nearly all experiments, as we know the conditions we use for induction are different from baseline and different from each other from previous work.

## Supporting information

Fasta file for all plasmid sequences

Complete PlcH Screen Data

Primers used in this study

## Acknowledgements

We would like to thank George O’Toole (Geisel School of Medicine at Dartmouth College) for the *rmcA* deletion and complementation strains.

