## Supplement Figures 1-3 for "A genome-wide screen in *Pseudomonas aeruginosa* identifies genes impacting production of the hemolytic phospholipase C/sphingomyelinase, PlcH"

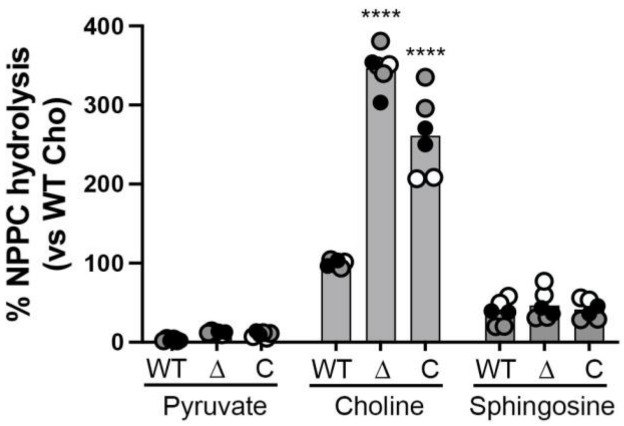


**Supplemental Figure 1. The effects of *cerN* deletion on PlcH expression.** PlcH activity as measured by NPPC hydrolysis comparing WT, Δ*cerN* (Δ), and the Δ*cerN attTn7::cerN* complementation strain (C) in the uninduced pyruvate condition versus induction with choline or induction with sphingosine. Statistical analysis for both panels done with two-way ANOVA with Dunnett's post-test with WT as the comparator strain in each condition (****, p<0.0001). All data points are shown and are colored by experiment with white circles for all replicates from experiment #1, grey from experiment #2, and black from experiment #3, and the overall means are represented by the bars. Only the means from each experiment are used in the statistical analyses for these panels (i.e. n = 3 per condition).

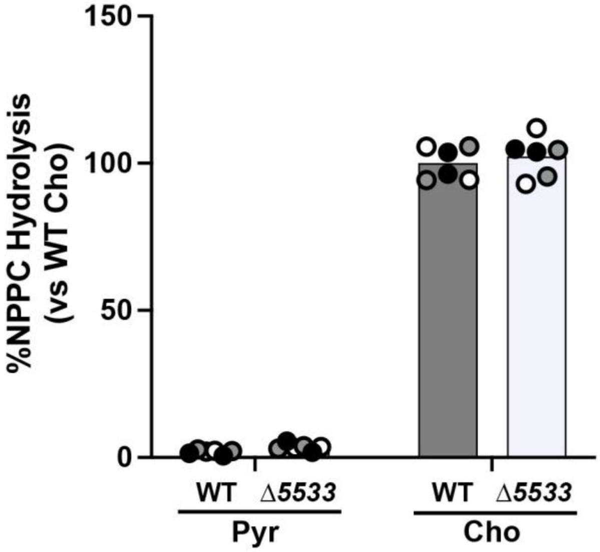


**Supplemental Figure 2. The effect of *PA5533* deletion on PlcH expression.** PlcH activity measured by NPPC hydrolysis comparing WT to the ΔPA5533 (Δ5533) deletion strain in the uninduced pyruvate condition (Pyr) versus the induced choline condition (Cho). There was no statistical difference between mutant and WT in either condition using two-way ANOVA with Dunnett's post-test comparing WT to the *PA5533* deletion strain in each condition. All data points are shown and are colored by experiment with white circles for all replicates from experiment #1, grey from experiment #2, and black from experiment #3, and the overall means are represented by the bars. Only the means from each experiment are used in the statistical analyses for these panels (i.e. n = 3 per condition).

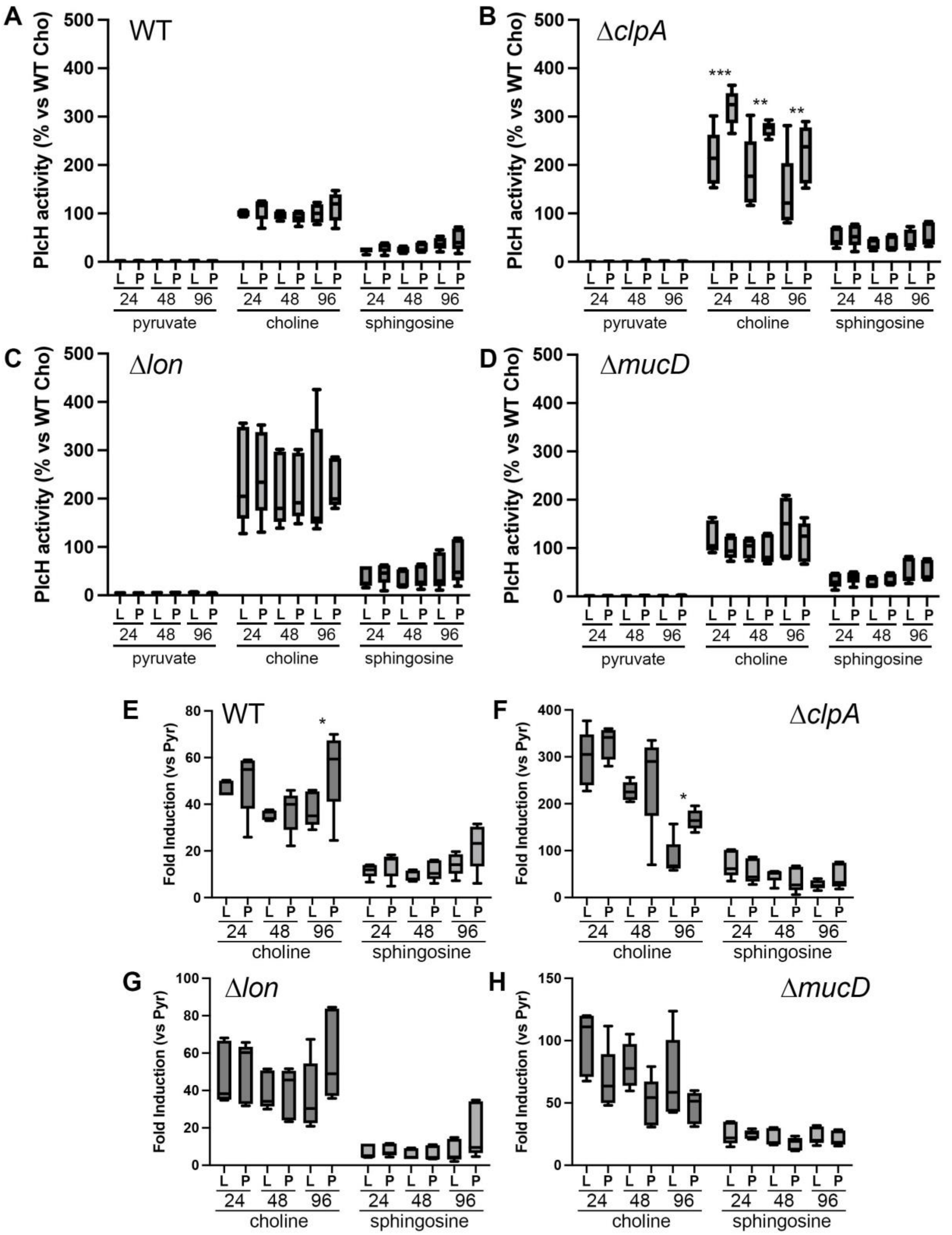


**Supplemental Figure 3. PlcH expression in WT vs *clpA*, *lon*, and *mucD* deletions in 24, 48, and 96-well formats and initiated from LB or PIA -80°C recovery plates.** PlcH activity measured by NPPC hydrolysis (**A-D**) and the fold-change versus pyruvate (**E-H**). The recovery plate media is noted by the L (LB) or P (PIA) immediately beneath each bar. The 24, 48, and 96 refer to the multiplate format and thus, well size. The data are grouped within each panel by inducing condition. The strain for is labeled in the top left or top right of each panel. Statistical analysis for each panel by two-way ANOVA followed by Tukey's post-test comparing all groups within each induction condition (*, $p<0.05$; **, $p<0.01$; ***, $p<0.001$). Only the significant changes between LB and PIA are noted, though there are some obvious changes comparing well sizes, which was of less interest.
